# Activation of a nerve injury transcriptional signature in airway-innervating sensory neurons after LPS induced lung inflammation

**DOI:** 10.1101/669374

**Authors:** Melanie Maya Kaelberer, Ana Isabel Caceres, Sven-Eric Jordt

## Abstract

The lungs, the immune and nervous systems functionally interact to respond to respiratory environmental exposures and infections. The lungs are innervated by vagal sensory neurons of the jugular and nodose ganglia, fused together in smaller mammals as the jugular-nodose complex (JNC). While the JNC shares properties with the other sensory ganglia, the trigeminal (TG) and dorsal root ganglia (DRG), these sensory structures express differential sets of genes that reflect their unique functionalities. Here, we used RNAseq in mice to identify the differential transcriptomes of the three sensory ganglia types. Using a fluorescent retrograde tracer and fluorescence-activated cell sorting we isolated a defined population of airway-innervating JNC neurons and determined their differential transcriptional map after pulmonary exposure to lipopolysaccharide (LPS), a major mediator of acute lung injury (ALI) and acute respiratory distress syndrome (ARDS) after infection with Gram-negative bacteria or inhalation of organic dust. JNC neurons activated an injury response program leading to increased expression of gene products such as the G-protein coupled receptors, Cckbr, inducing functional changes in neuronal sensitivity to peptides, and Gpr151, also rapidly induced upon neuropathic nerve injury in pain models. Unique JNC-specific transcripts, present at only minimal levels in TG, DRG and other organs, were identified. These included TMC3, encoding for a putative mechanosensor, and Urotensin 2B, a hypertensive peptide. These findings highlight the unique properties of the JNC and reveal that ALI/ARDS rapidly induce a nerve-injury related state changing vagal excitability.

**SIGNIFICANCE STATEMENT:** The lungs are innervated by sensory neurons of the jugular-nodose ganglia complex (JNC) that detect toxic exposures and interact with lung-resident cells and the immune system to respond to pathogens and inflammation. Here we report the expression of specific genes that differentiate these neurons from neurons in the other sensory ganglia, the trigeminal (TG) and dorsal root ganglia (DRG). Through nerve tracing we identified and isolated airway innervating JNC neurons and determined their differential transcriptional map after lung inflammation induced by a bacterial product, lipopolysaccharide (LPS). We observed the rapid activation of a nerve injury transcriptional program that increased nerve sensitivity to inflammation. This mechanism may result in more permanent nerve injury associated with chronic cough and other respiratory complications.

## INTRODUCTION

Lung inflammation is often caused by exposure to infectious agents or environmental pro-inflammatory toxicants. The culprits include Gram-negative bacteria, which contain lipopolysaccharide (LPS) in their outer membrane, or environmental organic dust contaminated with LPS. Intranasal endotoxin (LPS) administration in mice has been extensively used to study the role of innate immune responses in the pathogenesis of ALI/ARDS in pneumonia (1). LPS stimulates Toll-Like Receptor (TLR) 4 on mammalian cells, resulting in an innate immune response (2). Structural motifs on bacterial membranes stimulating TLRs not only elicit inflammation, but also itch and pain caused by activation of peripheral sensory neurons (3, 4). The immune and nervous systems are functionally interconnected. This interaction is bi-directional. Activation of some nerve types innervating the airways promotes inflammation and elicits mucus secretion, cough, sneezing and bronchoconstriction (5, 6), while activation of other nerve types dampens inflammatory responses (7).

The lungs are innervated by a complex network of sensory neurons (8, 9). The cell bodies of these neurons are located in the neck in the jugular and nodose ganglia of the vagus nerve, of neural crest and placodal origin, respectively (10–12). In humans and large mammals these ganglia are anatomically separated and innervate the lungs as well as the heart, stomach, kidneys and intestines (13). In mice, the ganglia are fused, forming the jugular-nodose complex (JNC). Each mouse JNC contains approximately 2000 neurons tightly surrounded by a large population of satellite glia cells. While this vagal structure shares similarities with other sensory ganglia, such as the dorsal root ganglia (DRG), and the trigeminal ganglia (TG), there are unique differences based on the different vagal innervation of organs, chemical and physical modalities sensed and peripheral and central response mechanisms. Previous studies identified transcription factors essential for developmental specification of the JNC and sets of ion channels and G-protein coupled receptors (GPCRs) involved in chemical and mechanical sensing. Some of these, such as the Transient Receptor Potential (TRP) ion channels, TRPA1 and TRPV1, are essential to maintain inflammation in asthma and trigger reflex responses such as cough as direct targets of pro-inflammatory environmental exposures (14–17). However, the repertoire of ion channels, G-protein coupled receptors and kinases contributing to sensing, signaling and reflex control by this essential structure remains understudied. Only a fraction of JNC neurons innervate the airways and lungs, with repertoires of expressed genes and functionalities likely different from JNC neurons innervating other organs (18). Hence, it is necessary to implement methods that selectively identify pulmonary sensory neurons to investigate their specific responses following exposure to LPS.

In this study we generate mouse bulk transcriptome datasets of the three sensory ganglia types, JNC, DRG, and TG, to identify JNC-specific transcripts. We use a fluorescent retrograde tracer to label and isolate airway-innervating neurons of the JNC (19), followed by cell sorting and RNA sequencing, to identify sets of genes differentially regulated upon lung exposure to LPS. Neuronal functional analysis is used to determine whether the observed transcriptional changes impact neuronal signaling and excitability.

## RESULTS

### Identification of jugular-nodose complex specific genes

To determine the genes uniquely expressed in the JNC, we compared the JNC transcriptome with that of the other sensory ganglia, the lumbar DRG and the TG. Bulk transcriptome datasets were generated by RNAseq of whole dissected mouse ganglia, containing all cell types, in triplicate (n=3). We used next-generation sequencing (NGS, Illumina HiSeq 2500) to quantify mRNA expression. NGS utilizes parallel processing by creating cDNA read fragments where many reads can be recorded at the same time (20). Each sample yielded > 29 million, 50 bp, reads.

Principal component analysis (PCA) revealed that components one and two were sufficient to describe 42% and 23% of variability between the different ganglia (*Figure 1A*). We then performed a two-group comparison of each ganglia group to the other two groups and identified the genes that were enriched, or depleted. Transcripts were filtered by using the p-value, p ≤ 0.0001. Applying these criteria, we identified 337 unique genes selectively transcribed at high levels in DRG, 1754 genes in the TG, and 791 genes in the JNC. Gene expression differences are displayed as heatmaps in *Figure 1B*. A full list of these genes with their transcript FPKM numbers are shown in *Data S1*.

**Figure 1:**
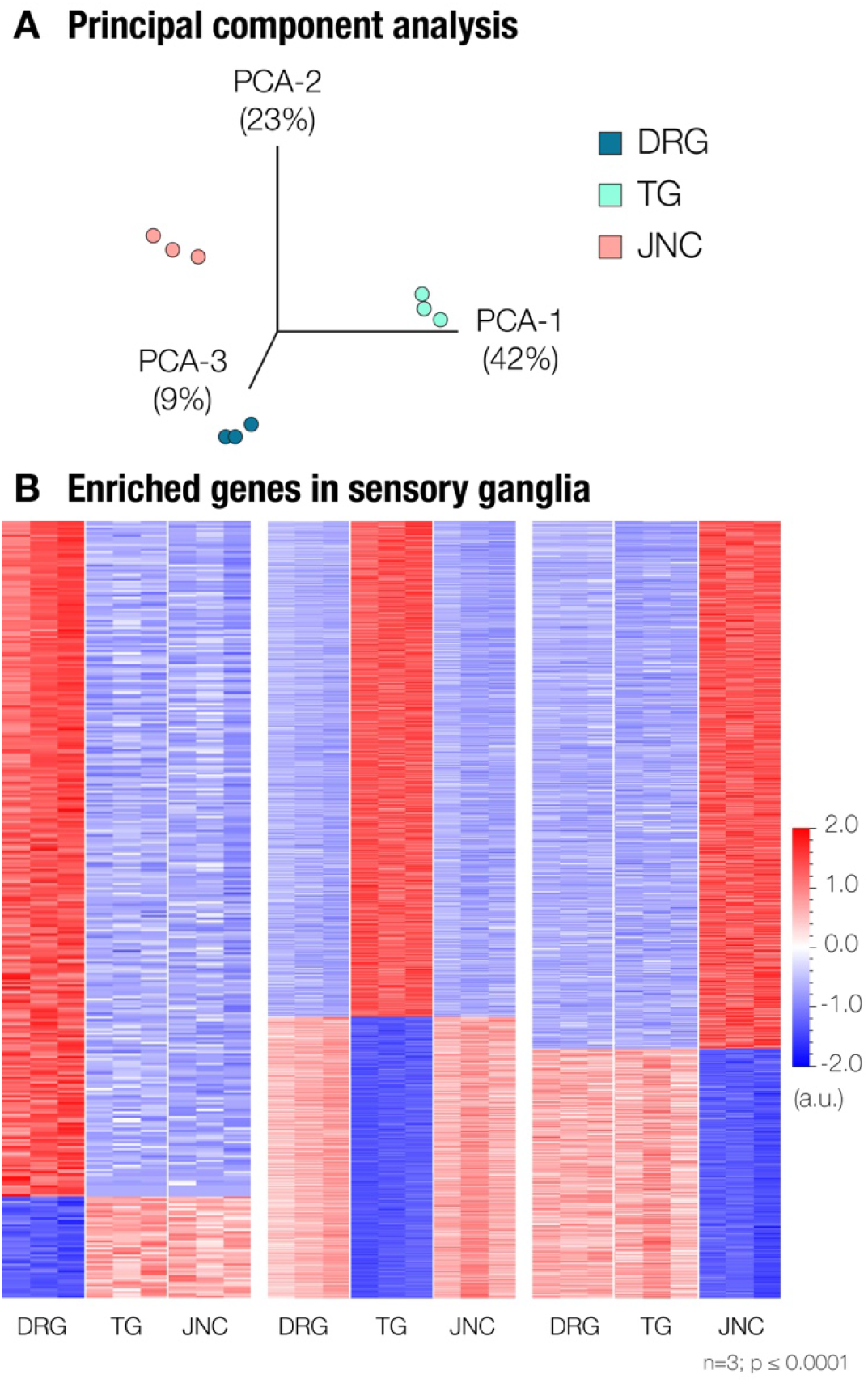
Transcriptional analysis of sensory ganglia. **(A)** Principal component analysis shows three groups associated with different sensory ganglia. **(B)** Heatmaps show a group of genes that are enriched in DRG (337 genes - *left*), TG (1754 genes - *center*), and JNC (791 genes - *right*). n=3, p ≤ 0.0001.

### Genes selectively expressed in the jugular-nodose complex

Next, we identified highly expressed genes in the JNC which had little to no expression in the DRG and TG *(Table 1)*. Leading the list were two known JNC marker genes, *Phox2b* and *P2rx2*. *Phox2b* encodes for a unique transcription factor specifying the development of the vagal ganglia (21, 22). Expression of *P2rx2,* encoding for a purinergic receptor ion channel, allows to distinguish between nodose and jugular neurons (23), which are fused into the JNC structure in the mouse. Previous studies confirm the lack of expression of these two genes in DRG and TG (24). These findings validate our approach, demonstrating that bulk sequencing of whole ganglia provides sufficient resolution to identify ganglia-specific selective gene expression.

**Table 1:**
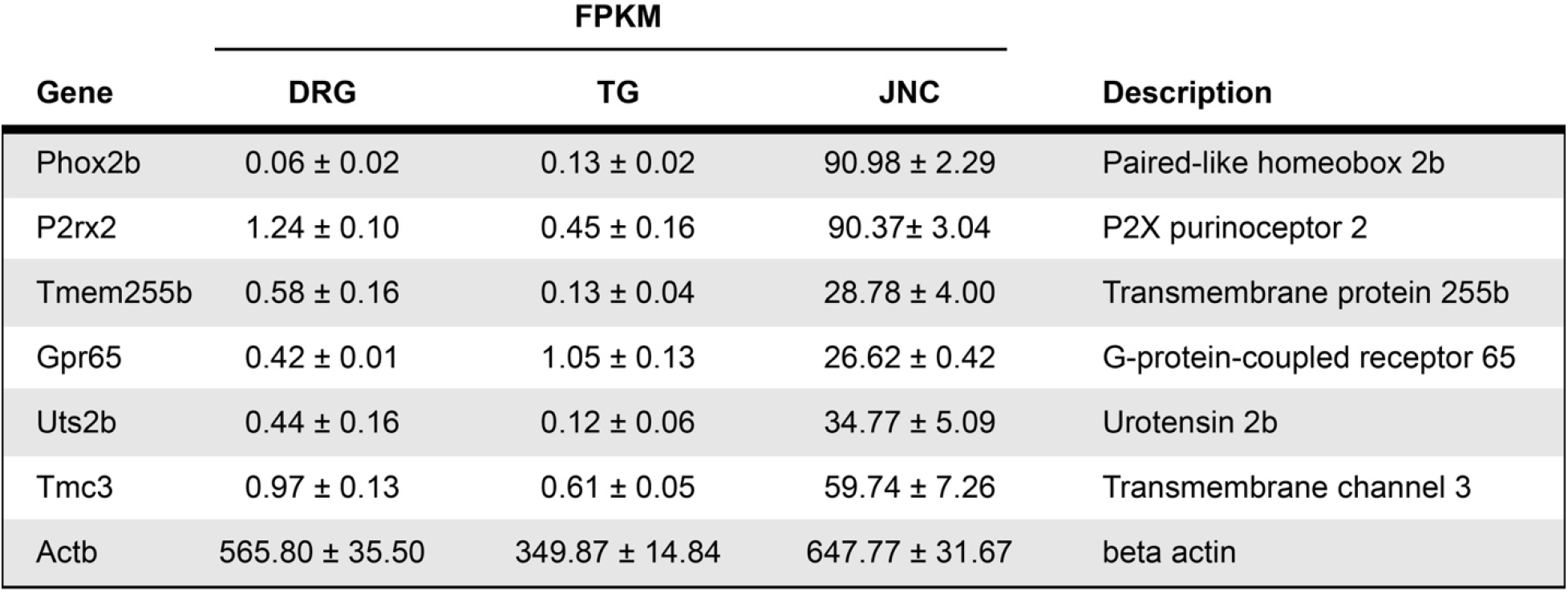
Enriched genes in the JNC. These genes are highly expressed in the JNC, but have very little to no expression in DRG and TG. Genes were identified as having a high FPKM in the JNC, and FPKMs < 2 in both the DRG and TG. *Actb* is included as a measure of total RNA. Values are reported as Mean ± S.E.M.

| Gene | FPKM |  |  | Description |
| --- | --- | --- | --- | --- |
|  | DRG | TG | JNC |  |
| Phox2b | 0.06 ± 0.02 | 0.13 ± 0.02 | 90.98 ± 2.29 | Paired-like homeobox 2b |
| P2rx2 | 1.24 ± 0.10 | 0.45 ± 0.16 | 90.37 ± 3.04 | P2X purinoceptor 2 |
| Tmem255b | 0.58 ± 0.16 | 0.13 ± 0.04 | 28.78 ± 4.00 | Transmembrane protein 255b |
| Gpr65 | 0.42 ± 0.01 | 1.05 ± 0.13 | 26.62 ± 0.42 | G-protein-coupled receptor 65 |
| Uts2b | 0.44 ± 0.16 | 0.12 ± 0.06 | 34.77 ± 5.09 | Urotensin 2b |
| Tmc3 | 0.97 ± 0.13 | 0.61 ± 0.05 | 59.74 ± 7.26 | Transmembrane channel 3 |
| Actb | 565.80 ± 35.50 | 349.87 ± 14.84 | 647.77 ± 31.67 | beta actin |

We then used real-time qPCR for validation of JNC-specific expression of the genes identified by RNAseq, and to compare expression in other neuronal tissues and major organs (*Figure 2*). *Tmem255b* (*Figure 2A*) and *Gpr65 (data not shown)*, were found to also be expressed in other tissues. However, *Uts2b* and *Tmc3* (*Figure 2B-C*) were found to be selectively expressed in only JNC, and not any other ganglia or organs, with *Tmc3* being particularly highly expressed (2714.50 ± 485.41 FPKM). The *Tmc3* gene belongs to the *Tmc* (Transmembrane Channel Like) family of genes, with eight genes known in the human and rodent genomes (25). We used qPCR to further examine the expression of all eight *Tmc* genes in sensory ganglia and other tissues (*Figure S1*). *Tmc3* was the only *Tmc* gene selectively expressed in the JNC, whereas *Tmc6* was highly expressed in all three types of sensory ganglia.

**Figure 2:**
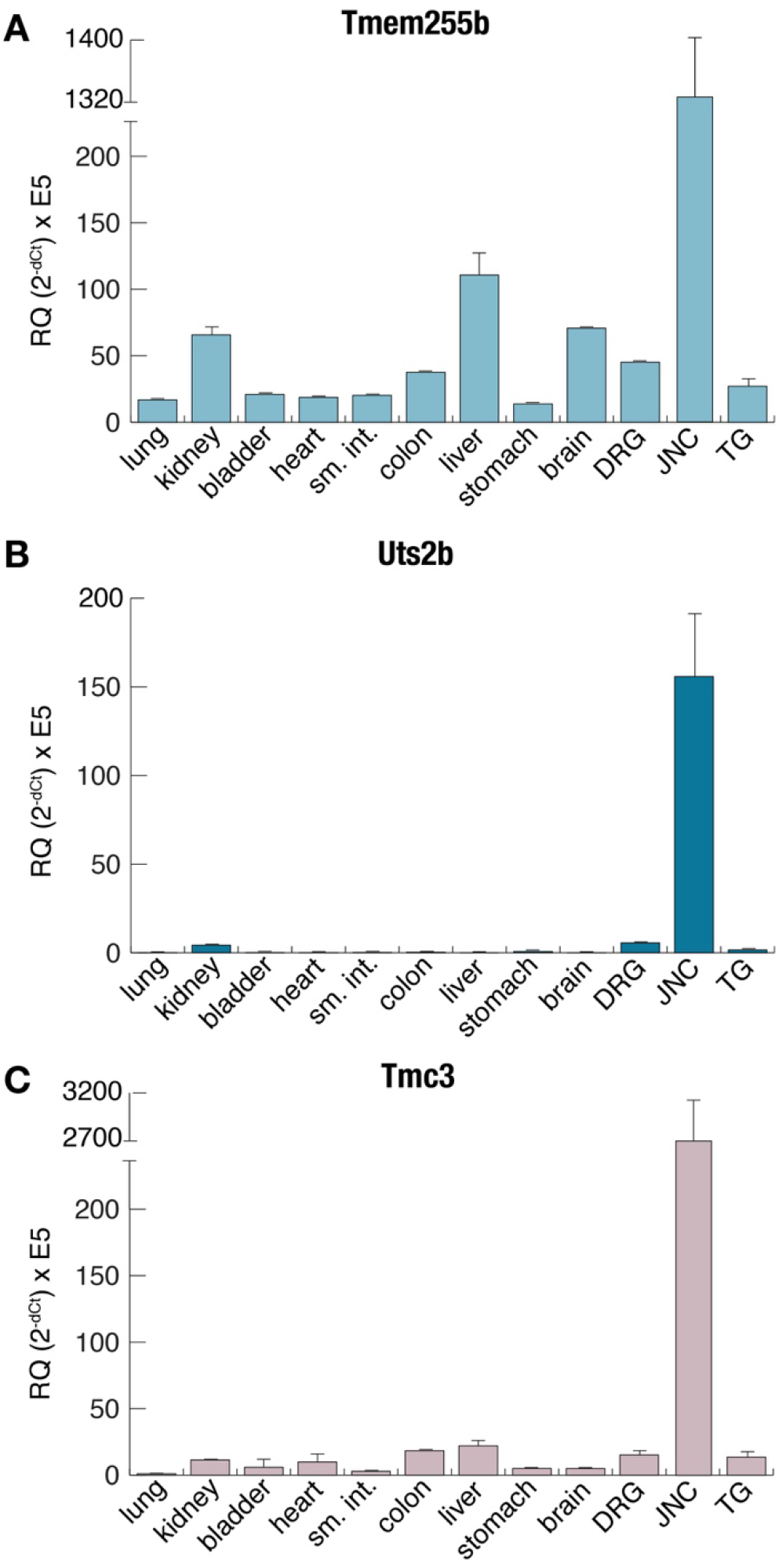
Tissue qPCR expression of JNC highly expressed genes. Real time qPCR was used to determine the expression of **(A)** *Tmem255b*, **(B)** *Uts2b*, and **(C)** *Tmc3* in different tissues. *Actb* was used as the housekeeping gene, n=4, values calculated as 2-dCt x E5, mean ± S.E.M.

Less stringent selectivity criteria identified additional genes with enriched expression in the JNC. These included *Chrna3* and *Chrnb4*, encoding for nicotinic receptor sub-units, with DRG and TG showing some baseline expression (*Figure S2*).

### Transcriptomic responses in whole JNC and retrogradely traced lung-innervating JNC neurons to Lipopolysaccharide-induced pulmonary inflammation

For induction of pulmonary inflammation, mice were given intranasal instillations of 8 µg LPS in 40 µl saline, once daily on three consecutive days. The vehicle control group was given 40 µl of saline only following the same schedule. On the fourth day the bronchoalveolar lavage was performed to collect pulmonary cells, followed by immediate dissection of lungs and neuronal tissues. The retrograde tracer Fast Blue was used to label and identify JNC neurons innervating the lungs. 40 µl of 350 µM Fast Blue were administered intranasally, one day before the first LPS (or vehicle) administration (*Figure 3A*). Fast Blue labeled neurons were clearly identifiable in JNC cryosections and whole mount preparations (19) (*Figure 3A; Movie S1*). One day after the 3^rd^ LPS administration JNC were dissected and sorted by FACS, into Fast Blue positive (lung innervating) or negative pools. Sorted cell pools were processed for RNAseq. Each sample had 29-30 million successful reads, and greater than 80% read alignment (*Table S1*). In both Fast Blue positive and Fast Blue negative populations the highly enriched JNC genes (*Table S2*) were similarly expressed. Further supporting that the enriched genes we describe are global JNC markers.

**Figure 3:**
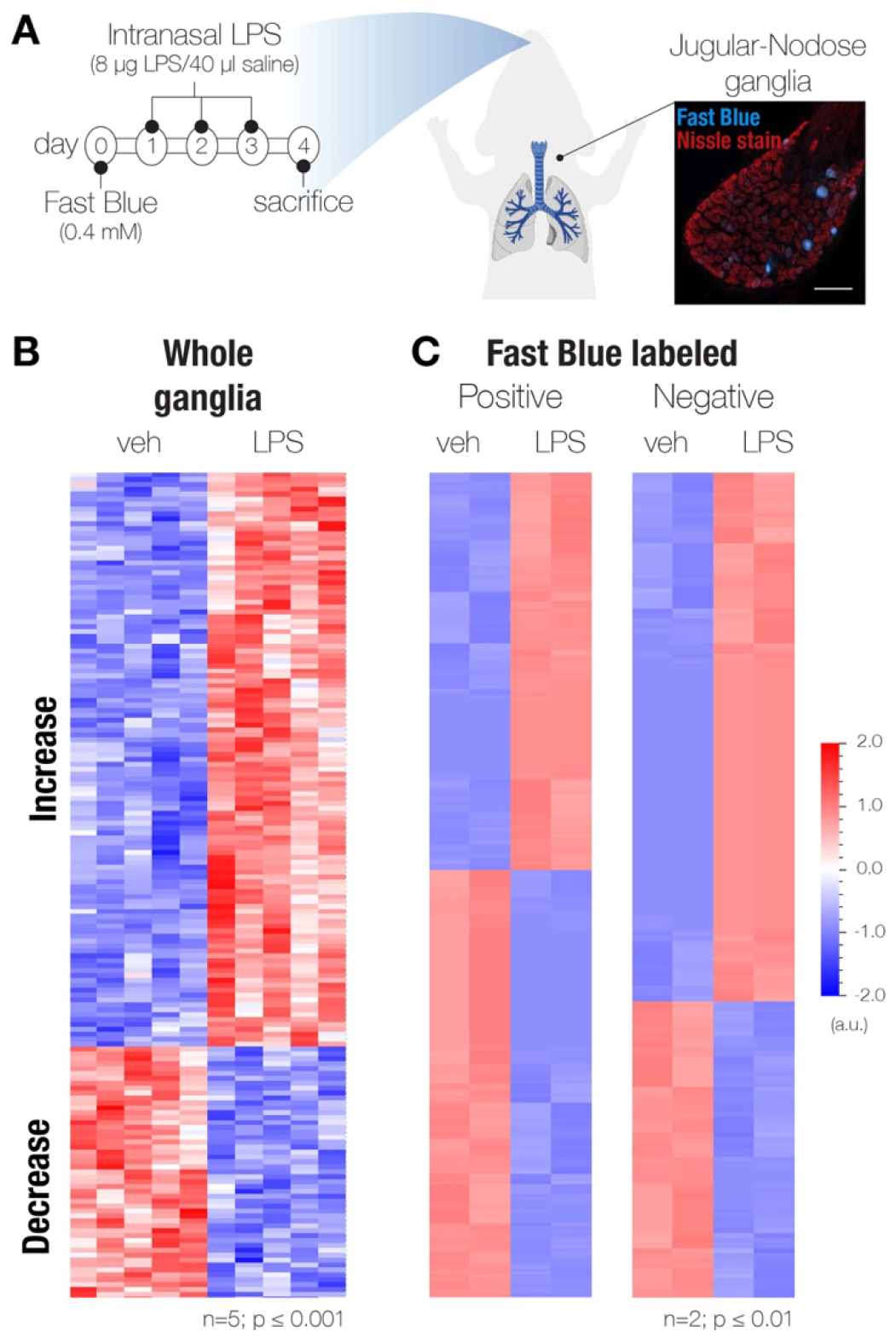
Lipopolysaccharide (LPS) lung exposure changes transcriptional profile of airway innervating JNC neurons. **(A)** Timeline for exposure to Fast Blue tracing dye and LPS (8µg/40 µl saline), vehicle mice were just given saline. **(B)** Heatmap showing 169 genes that are enriched in whole JNC ganglia with and without LPS-induced inflammation. n=5, p ≤ 0.001. **(C)** Heatmap of genes enriched in sorted JNC neurons positive for Fast Blue (255 genes - *left*) or negative for Fast Blue (164 genes - *right*); n=2, p ≤ 0.01.

In addition to the Fast Blue-positive and -negative cells, we also created transcriptome datasets for whole, un-dissociated ganglia from LPS- and vehicle-treated mice. These datasets contain information from all ganglia cell types, both neurons and supporting satellite cells, including immune cell populations located within the JNC. Due the restriction of Fast Blue to neuronal cells, the Fast Blue-positive population represents the pure lung-innervating population of JNC neurons, while the Fast Blue-negative population contains both the non lung-innervating JNC neurons and satellite cells. Heat maps illustrate the populations of up- and down-regulated genes in whole ganglia and the sorted Fast Blue-positive and –negative cell populations (*Figure 3B, C*). The full list of genes with expression altered following LPS administration and their FPKM values can be found in *Data S2*. A full list of genes differentially regulated in lung-innervating JNC neurons was generated by comparing transcriptome datasets from Fast Blue positive and negative populations of mice treated with vehicle or LPS (*Data S3*). Figure 4 ranks the most strongly regulated genes found in the whole ganglia (*Figure 4A*), and in both Fast Blue positive *(Figure 4B*), and Fast Blue negative populations (*Figure 4C*). Genes that are significantly upregulated in the whole ganglia transcriptome dataset, and also upregulated in the Fast Blue negative population but have an FPKM value of 0 in the Fast Blue-positive population most likely indicate of expression changes in the satellite cells, immune cells, or neurons that innervate other tissue (*Figure 4A*). These genes include *Lrg1, Lcn2, Selp, Gbp4, Xdh,* and *Gbp5.* Of the upregulated genes in the whole ganglia *Lrg1, Lcn2, Gbp2, Chi3l3,* and *Ch25h* were shown to also be upregulated locally in lung tissue after LPS exposure (26). These genes are likely associated with the innate immune response and may be regulated in immune cells in both the lungs and in JNC.

**Figure 4:**
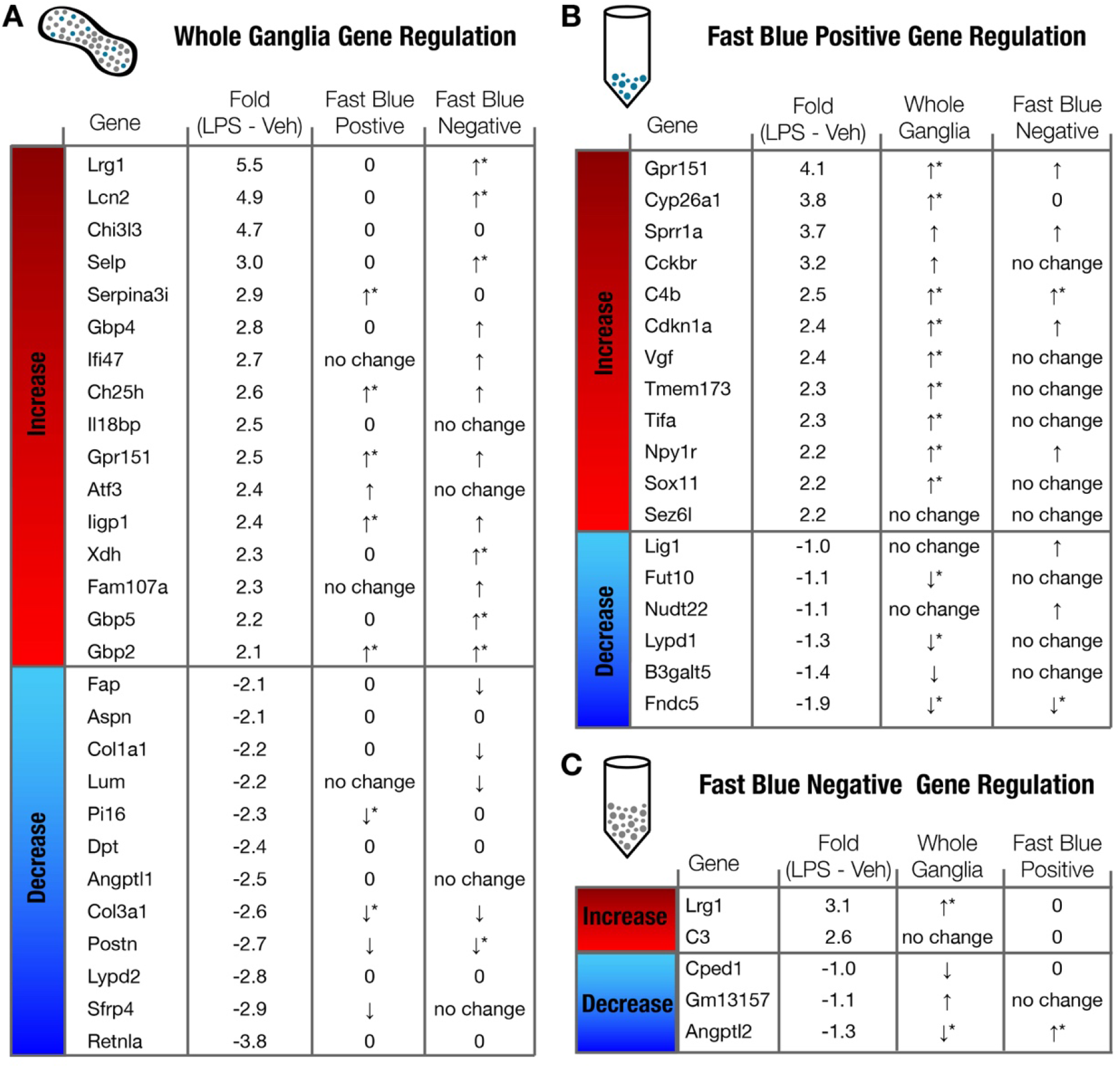
LPS-induced gene regulation. Fold changes are show as base two for LPS – vehicle (Veh) [fold-change = log_2_(LPS FPKM) – log_2_(Veh FPKM)]. **(A)** Whole ganglia gene regulation fold changes compared Fast Blue positive and negative sequencing data. **(B)** Fast Blue positive gene regulation fold changes compared to whole ganglia and Fast Blue negative data. **(C)** Fast Blue negative gene regulation fold changes compared to whole ganglia and Fast Blue positive data. No change indicates a significance value of p > 0.2, an arrow (no star) indicates 0.2 < p < 0.05 and an arrow with a start indicates p < 0.05. A “0” means FPKM < 1.

LPS lung exposure only affected transcription of very few genes in the Fast Blue negative (non-lung-innervating) population (*Figure 4C*). This is an important observation suggesting that JNC nerve populations respond selectively to changes in the pathophysiological states of their target organ, while neighboring populations innervating other organs remain unaffected. The two genes upregulated in the FB-population, *Lrg1* and *C3,* are both associated with innate immunity (27, 28). The selective regulation of these genes suggests that either immune cells are maturing in the JNC (29), or satellite cells are taking on immune cell characteristics (30, 31). Satellite glia cell activation may help maintain neuronal changes during inflammation. Genes upregulated in the Fast Blue positive population of lung-innervating JNC neurons (*Figure 4B*) include *Gpr151, Cckbr*, both encoding for G-protein coupled receptors (GPCRs) and *Sprr1a, Sox11, Sez6l, Vgf,* and the transcription factor *Atf3* (32–34).

Changes in transcript levels of the above genes in the JNC following LPS exposure were validated by real time qPCR and compared with levels in cDNA generated from lumbar DRG of the same animals (*Figure 5A*). This analysis confirmed their strong upregulation in JNC, but only minimal expression changes in DRG (*Figure 5A*). Two genes, *Atf3* and *Gpr151*, were moderately upregulated in DRG compared to controls, suggesting that these are highly sensitive to beginning systemic changes.

**Figure 5:**
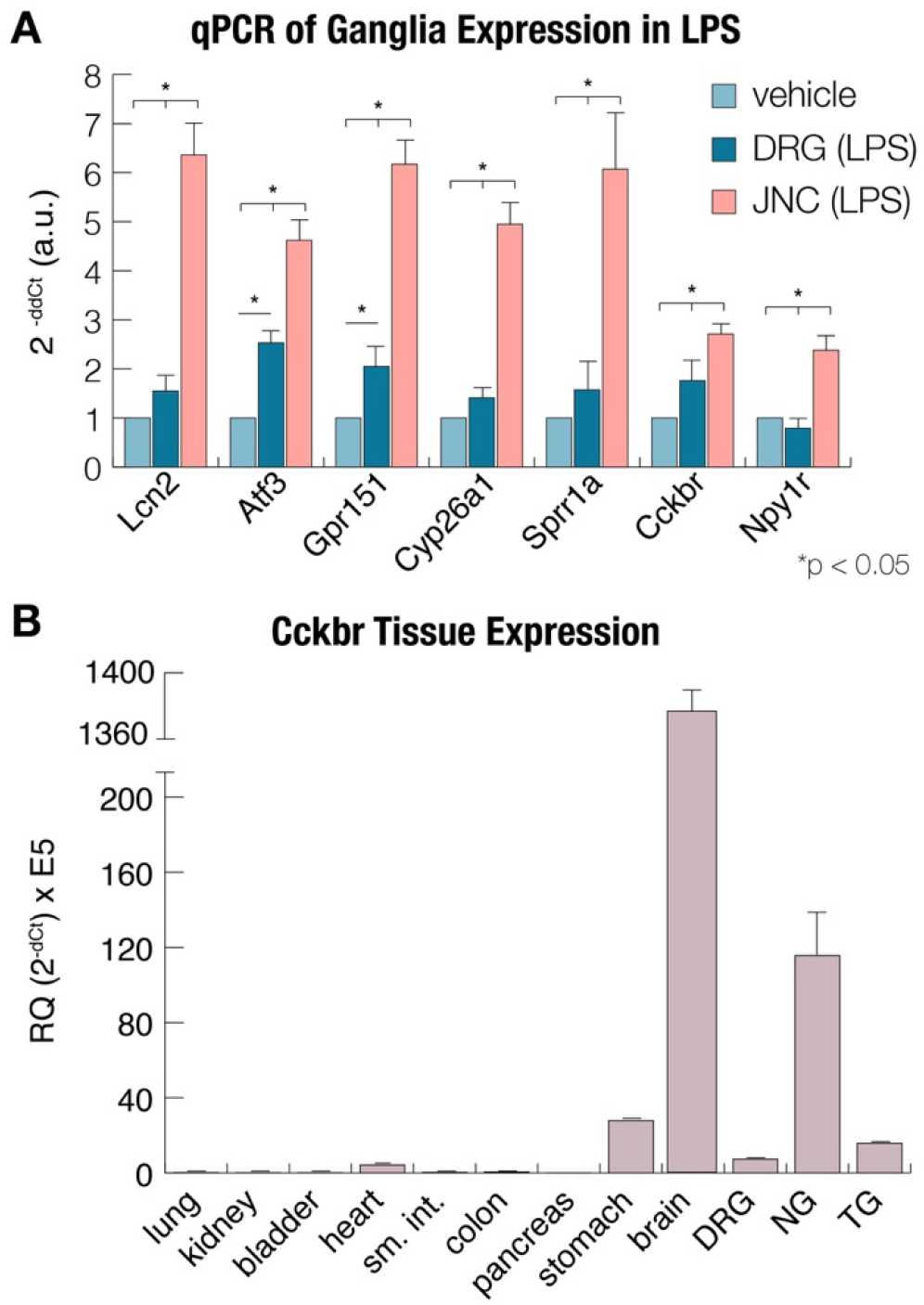
Real time qPCR confirmation of gene expression. **(A)** Real time qPCR was used to validate RNA sequencing findings. The ΔΔCt measurement was used to calculate fold change of each gene compared to vehicle. **(B)** Tissue specificity of *Cckbr* expression shows that the JNC is enriched for the gastrin/CCK receptor. For each group n = 4-6 and *Actb* was used as the housekeeping gene. Values are reported as mean ± SEM.

### Correlation of transcriptome and functional changes in JNC neurons after LPS treatment

The gene encoding for the cholecystokinin B receptor (CCKBR), *Cckbr*, was among the most strongly LPS-induced genes in the JNC. While the majority of the identified LPS-induced gene products are orphan receptors or not targetable, highly selective agonists of CCKBR are available to examine whether the observed changes in JNC transcriptome levels resulted in neuronal functional changes. Real time qPCR analysis was performed to compare Cckbr transcript levels in JNC to those in other sensory ganglia and other tissues (*Figure 5B*). *Cckbr* transcript levels are highest in brain, followed by JNC. As shown previously, the stomach shows moderate expression (35–37). *Cckbr* transcript levels were higher in JNC compared to DRG and TG (*Figure 5B*).

To examine whether LPS exposure-induced *Cckbr* transcription resulted in functional changes, JNC neurons were dissociated one day after the last of three pulmonary LPS treatments. Neurons were analyzed by fluorescent calcium imaging 2 hours after plating. Gastrin (1 nM), a CCKBR-specific agonist, was used for stimulation following recording of baselines (38). Subsequently, JNC neurons were superfused with capsaicin (100 nM) to excite TRPV1-expressing neurons (C-fibers), followed by KCl (30 mM) as a strong depolarizing stimulus identifying all neurons (*Figure 6A*).

**Figure 6:**
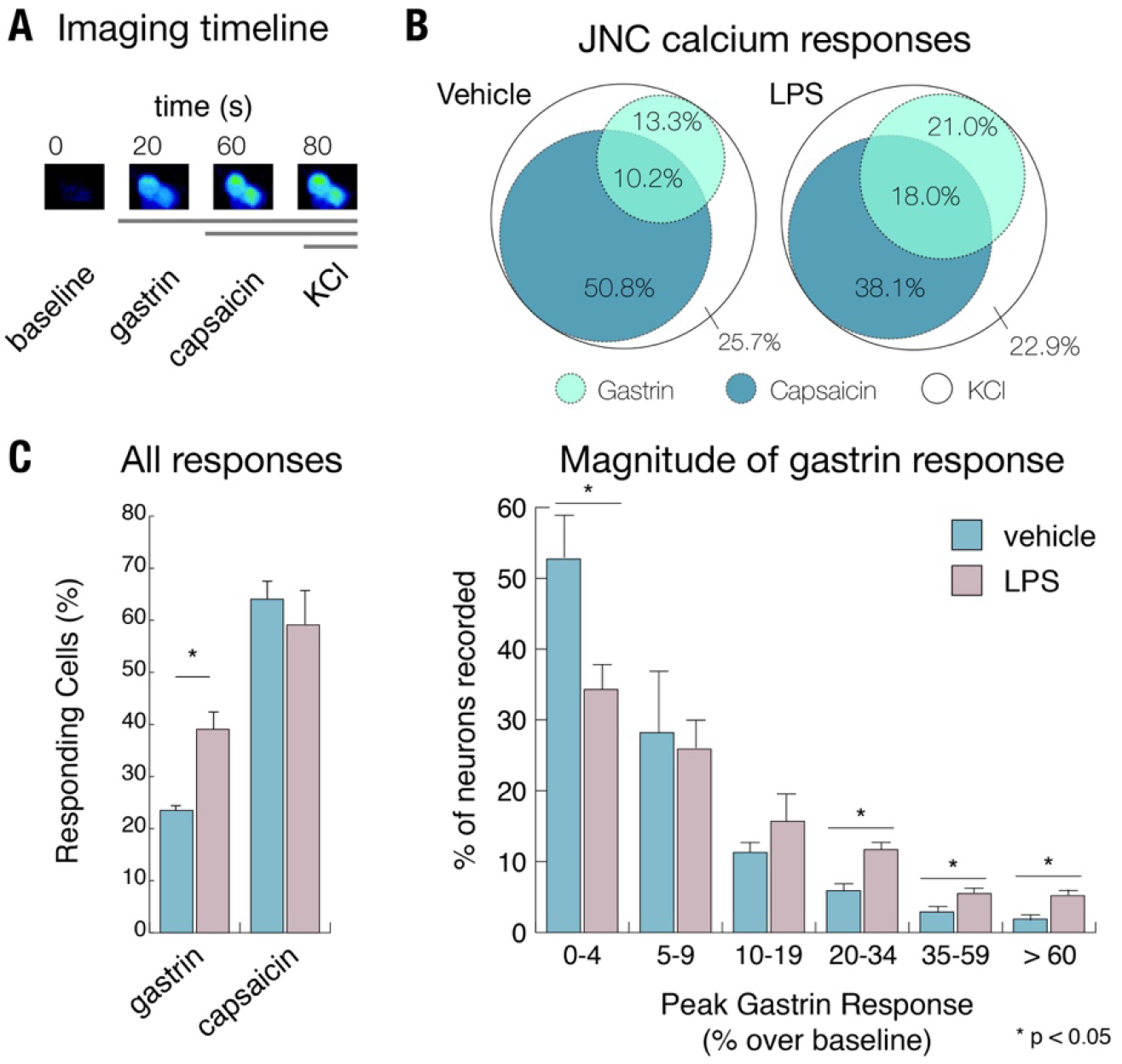
CCKBR neuronal activity increases after LPS-induced lung inflammation. **(A)** Imaging timeline for stimuli (gastrin - 1nM, capsaicin - 100nM, and KCl - 30mM) application. **(B)** A Venn diagram showing the overlap between the gastrin and capsaicin responding cells. **(C)** *Left* - The percent of cells that responded to gastrin in the vehicle (healthy) versus LPS (sick) mice significantly increased. *Right* - The magnitude of the gastrin response over baseline in vehicle (healthy) versus LPS (sick) neurons also significantly increased. The capsaicin response remained constant after treatment. The total number of neurons recorded was 1,542 vehicle and 1,270 LPS neurons. A positive calcium response was defined as a >10% increase, over baseline, of the ratiometric signal (F340/380). N=5 mice for each group. Data are reported as mean ± SEM.

JNC neurons dissociated from LPS-treated animals contained larger numbers of neurons responding to gastrin with calcium influx than neurons from vehicle-treated animals. The percentage of gastrin-responding cells significantly increased from 23.5% ± 1.2 in the vehicle group to 39.0% ± 4.8 in the LPS-treated group (*Figure 6B-C*). The number of capsaicin-sensitive neurons was not significantly different between groups. When we plotted the peak amplitude of the gastrin response based on the number of neurons recorded from (*Figure 6C-right*), we saw a significant increase in the larger calcium responses (> 20% increase over baseline) and a decrease in the number of neurons that did not respond (< 4% increase over baseline). This implies that more neurons express functional CCKB receptor, and are responding more strongly, after LPS-induced lung inflammation.

## DISCUSSION

Sensory ganglia innervate different regions of the body and different organs. As such, their molecular profiles are expected to differ based on target organs and functional needs. Our comparison of sensory ganglia transcriptome datasets revealed specialized molecular markers expressed in the murine vagal sensory neurons of the JNC. We identified several uniquely expressed transcripts in JNC neurons:

### Tmc3

Tmc3 is a gene of the Tmc (Transmembrane Channel like) family of eight mammalian genes (39). While the function of TMC3 in JNC neurons is not known, TMC1 and TMC2 were identified as potential components of the auditory hair cell mechanotransduction complex. Mutations in Tmc1 and Tmc2 were linked to deafness in multiple human populations (25, 40). These findings suggest that some Tmc proteins may contribute to mechanical sensing in other sensory systems. In *C. elegans*, tmc-1 was shown to be a sodium-sensitive ion channel, implicating a role in homeostatic ion sensing or, potentially, osmotic sensing (41). A longitudinal genome-wide study linked a polymorphism near the IL6/STARD5/TMC3 locus on chromosome 15 to a higher rate of lung function decline in humans (42). However, it is not clear which of the three genes in this locus are differentially regulated due to the polymorphisms. The highly selective expression of Tmc3 in JNC, and not in other sensory ganglia or other organs, warrants further studies examining its role in JNC development, target organ innervation and sensory mechanisms.

### Uts2b

Similar to Tmc3, the gene Uts2b proved to be selectively expressed in the JNC, with minimal expression in TG, DRG and kidney, but undetectable in other major organs. Uts2b encodes for the peptide Urotensin II, a potent vasoconstrictor involved in blood pressure regulation (43). Sub-cutaneous injection of urotensin-II in monkeys caused cardiac dysfunction (44). Receptors for Urotensin II were targeted for drug development with the aim to generate antihypertensive treatments. Some neuronal expression of Uts2b was reported in in motor neurons of the spinal cord (45). The kidney was discussed as the major systemic source of Urotensin II (46, 47). Our data suggest that, in fact, Urotensin II is a vagal sensory neuropeptide, potentially released by JNC neurons peripherally upon vagal nerve stimulation. This finding is supported by recent single cell transcriptome analysis of vagal sensory nerves, identifying an Utsb2b-positive neuronal population hypothesized to contribute to digestive tract innervation (10). Uts2b transcript levels were low in airway-innervating JNC neurons in our study, suggesting a minor role in control of respiratory function (*Figure S3*). A Urotensin II antagonist has been tested as an oral treatment in human asthmatic patients but failed to produce positive bronchodilation effects (48).

### Tmem255b

Expression of Tmem255b was confirmed to be high in the JNC, however it also has basal low-level expression in all other tissues we tested. Not much is known about TMEM255B. In humans, Tmem255b expression is increased in colon cancer (49) and a mutation was found in an obese child (50). These findings do not establish a functional role or pattern for this gene encoding for a hypothetical membrane protein. Its function in vagal development and sensing has yet to be determined.

### Gpr65

The G-protein-coupled receptor Gpr65 was originally identified as a receptor for certain lipid species and as a proton receptor activated by acidity (51–53). While expression of Gpr65 in JNC neurons was reported previously (23) the authors did not observe any respiratory effects of Gpr65 gene ablation in mice. This is consistent with our data that did not detect significant Gpr65 expression in the airway-innervating population of JNC neurons. Instead, this gene likely to contributes to sensing and control of other visceral organs. Indeed, a polymorphism associated with the Gpr65 gene was shown to determine the severity of inflammatory bowel disease (IBD) (54).

Use of the retrograde fluorescent tracer, Fast Blue, enabled us to identify a population of JNC sensory neurons that innervates the airways and lungs. This approach may not have labelled all lung-innervating sensory nerves since some JNC neurons may not extend to airway surfaces within the lung where Fast Blue uptake occurs after aspiration. Nevertheless, a substantial number of neurons was labeled enabling us to gather transcriptome data from pooled Fast Blue-positive neurons and to identify LPS-induced genes. The transcriptome dataset is most probably from a heterogeneous lung-innervating neuronal population and needs to be further analyzed using single cell approaches that so far have not used organ-specific retrograde tracing methods (10). Our approach was sensitive enough to identify the major LPS-induced neuronal genes and to detect essential differences between datasets produced from other whole sensory ganglia (TG, DRG). The following genes were induced following LPS-exposure, with information provided about their expression site (lung-innervating nerves, nerves innervating other organs, satellite cells) and potential functions:

### Gpr151

This gene is the most strongly LPS-induced gene in lung-innervating JNC neurons, encoding for a G-protein-coupled receptor that was first described as a galanin receptor like receptor (GALRL) in 2004. However, this receptor is sensitive to galanin only concentrations by far exceeding physiological levels. GPR151 may have other endogenous agonists that remain to be identified (55). Intriguingly Gpr151 is strongly induced in DRG after neuropathic injury in pain models, described in several recent studies (32, 56–58). Similar to GPR65, GPR151 was shown to be activated by acidity; however, its role in pain signaling may be stimulus-dependent (32, 56–58). The induction of this gene in JNC after LPS exposure suggest that JNC neurons experience neuropathic injury and activate a response program with partial similarities to injury-induced gene sets in DRG neurons.

### Lcn2

The gene Lcn2 encodes the protein lipocalin-2, which is associated with innate immune responses to bacteria activated downstream of TLR4 activation by LPS. Lipocalin-2 is found in neutrophils as well as endothelial cells, astrocytes and microglia (30). In the brain and spinal cord inflammation activates microglia. Once activated, they secrete LCN2 which is toxic to neurons (59, 60). Upregulation of the Lcn2 gene following LPS exposure was observed in the bulk whole ganglia datasets, but not in the Fast Blue positive (lung-innervating) neurons, which implies this gene is induced by LPS in the satellite cells, or neurons that innervate other tissue.

### Atf3/Sprr1a

Activated transcription factor 3 (ATF3) plays multiple roles in gene expression regulation in different cell types. In Atf3−/− mice, neutrophils are unresponsive to chemotaxic gradients and unable to find their target (24). In DRG sensory neurons however, Atf3 expression is associated with nerve injury and promotes nerve growth and regeneration by turning on Sprr1a expression (25, 26). SPRR1A associates with F-actin to promote nerve growth (61). This activated pathway in neurons may promote nerve growth and repair of JNC neurons after LPS exposure.

### Cyp26a1

CYP26A1 is a cytochrome enzyme that breaks down retinoic acid, preventing excessive retinoic acid signaling. Retinoic acid plays an important role in cell proliferation neuronal and limb development (62). The Cyp26a1−/− mouse is embryonic lethal (63). Activated microglia has also been shown to produce CYP26A1 (64). The role of Cyp26a1 expression in JNC neurons after LPS exposure is unclear, but may contribute to nerve regeneration or growth and sprouting following nerve injury.

### Npy1r

NPY-R1 is a receptor for Neuropeptide Y (NPY), a neuropeptide upregulated in the lungs during inflammation (65). Its receptor can be found on immune cells and neurons. In DRG neurons NPY-R1 is found mostly in pain-transducing neurons and its activation leads to an inhibitory current (66, 67). This presents a complex scenario in which some neurons may become sensitized by inflammation and other inhibited dure to increased sensitivity to NPY.

### Cckbr

Our unbiased approach revealed that the expression of Cckbr, encoding for the Cholecystokinin B Receptor, is upregulated in the JNC neurons that innervate the airways following LPS exposure. CCKBR, a G-protein coupled receptor, is the only known target of gastrin, a peptide produced in the pancreas and other digestive organs, first discovered in 1993 (38). CCKBR is also activated by cholecystokinin (CCK), a peptide implicated in pain signaling in neuropathic pain within the spinal cord and, potentially, in the periphery.

The analysis of nocifensive behavioral responses of Cckbr−/− mice in the chronic constriction model of nerve injury observed a lack of allodynia, demonstrating this gene fulfills a role in pain perception (68). Interestingly, the authors reported increased expression of TLR4 in the brain of Cckbr−/− mice compared to wild type (68). This suggests that Cckbr may regulate neuronal sensitivity to innate immune stimuli. Cckbr expression was also upregulated in DRG neurons in a burn injury model, and inhibition of CCKBR was shown to have analgesic effect (69).

While the expression of gastrin is restricted to digestive tissues, CCK peptides are expressed within the lung, by pulmonary neuroendocrine cells (PNECs) and other cell types (70–72). A CCK peptide was shown to induce bronchoconstriction in humans and guinea pigs (73). Exogenously administered CCK peptides were shown to alter cytokine expression and counteracted inflammatory responses in a LPS-induced mode rat model of endotoxic shock (74, 75). CCK signaling has also been implicated in lung injury due to pancreatitis CCKBR is also involved in proliferative responses such as cell growth in the stomach lining, and in aberrant proliferative responses pancreatic cancer cells (76), colon cancer (77), and small cell lung cancer (78). In the digestive tract, the CCK/gastrin/CCKBR system responds to bacterial infection by Helicobacter pylori (H. pylori), Gram-negative bacteria, initiating an immune response and increasing gastrin secretion (79).

To infer the function of the CCKBR receptor in neurons it is helpful to compare with the function of the other genes induced by LPS in the in Fast Blue positive neurons. Some of these genes, including Atf3 and Sprr1a, are also induced in DRG neurons following neuropathic injury in pain models, and may initiate either a nerve repair program, or stimulate pathological nerve branching and sprouting, potentially leading to increased sensitivity and chronic pain perception (58, 80). CCKBR may contribute to these mechanisms as it contributes to proliferative and growth mechanisms in other cell types. Whether LPS-induced increase in expression of these genes counteracts inflammatory responses, or contributes to transient or permanent neuropathic states increasing or dampening vagal reflex responses such as cough, glandular secretions and cardiovascular reflexes remains to be established (81–83). Some of the identified receptors and membrane proteins may serve as targets to interfere with exaggerated vagal responses to LPS and suppress pro-inflammatory mechanisms.

## MATERIALS AND METHODS

### Animals

C57BL/6 mice from Charles River Laboratories (Wilmington, MA) were used (female, 8-12 weeks old). Procedures were approved by the Institutional Animal Care and Use Committee (IACUC) of Duke University. Mice were housed in standard environmental conditions (12-hour light-dark cycle and 23 °C) at facilities accredited by the Association for Assessment and Accreditation of Laboratory Animal Care. Food and water were provided ad libitum. All the animal protocols were submitted to, and approved by, the Institutional Animal Care and Use Committees of Duke University. All the research with mice conform to the ARRIVE guidelines for animal studies (84).

### Fast Blue retrograde labeling

Under light inhaled anesthesia (sevoflurane, open-drop exposure method) mice received a 40 µl intranasal instillation of 350 µM Fast Blue (Polyscience, Inc) and 1% DMSO in saline. Mice were sacrificed 3-14 days after Fast Blue administration and the jugular-nodose complex were collected (19).

For visualization, JNC were fixed in 4% PFA overnight, and then immersed in 10%, 20% and 30% sucrose solutions. Tissue was embedded in OCT, sectioned using a Cryostat at 20µm sections, and collected on slides. Slides were post-fixed in 10%NBF, washed with PBS, and treated with NeuroTrace 530/615 red fluorescence Nissl stain (Thermo, cat#: N21482) according to the manufacturers’ recommendations. Sections were imaged using a fluorescence Zeiss microscope.

### Lipopolysaccharide airway exposure

Mice received three intranasal administrations with either lipopolysaccharide (*E. Coli*, 0111:B4 Calbiochem) or vehicle (saline). Lightly anesthetized mice (sevoflurane) received 8 µg of LPS in 40 µl of saline or saline alone on days 1, 2, and 3. The mice underwent bronchoalveolar lavage and were sacrificed for tissue collection 24 hours after the last LPS administration.

### Bronchoalveolar lavage (BAL) and leukocyte analysis

BAL was performed by cannulation of the trachea and gentle instillation/aspiration (3 times) of 1.0 ml PBS with 0.1% BSA and protease inhibitor cocktail tablets (Roche, Indianapolis, IN). Animals were then perfused with PBS before tissue collection. The lavage fluid was centrifuged and the supernatant was discarded. The cell pellet was treated with red blood cell lysing buffer (BD Biosciences, San Diego, CA), washed, and resuspended in 200 µl of PBS. Total cell counts were determined with a hemocytometer (Scil Vet ABC, Gurnee, IL), centrifuged onto cytoslides (Shandon Cytospin 3 cytocentrifuge, Thermo Electron Corporation, IL, USA), and stained with Diff-Quick (Dade-Behring Inc., Newark, DE). Differential cell counts were obtained using a microscope to count a minimum of 200 cells/slide using standard morphological and staining criteria.

### Dissociation of primary neurons

JNC, DRG and/or TG were removed from the animal and immediately placed into 1 mL neuronal media (Advanced DMEM/F12, Glutamax, penicillin/streptomycin, 1 M HEPES, N2 supplement, B27). The tissue was washed once with calcium free PBS and resuspended in digestion media (55 µg Liberase in 1 ml neuronal media). Tissue was digested at 37 °C for 45-60 minutes, washed, resuspended in 200 µl neuronal media and pipetted up and down with 200 µl tip until tissue was completely disrupted. Cells were passed through a 70 µm cell strainer then added to a Percoll Gradient (28% over 12.5%). For FAC sorting, dissociated cells were kept on ice. For calcium imaging, cells were plated in an 8-well chamber pre-coated with poly-D-lysine and laminin.

### FAC sorting

Flow cytometry was done on a BD FACS Aria II running Diva 8 software (BD Biosciences). For cell sorting, a large nozzle (100 µm) and low pressure (20 psi) was used. To set up the sorting gates, dissociated non-lung innervating dorsal root ganglia were used. The positive control was dissociated dorsal root ganglia incubated with 350 µM Fast Blue in vitro. Cells were sorted directly into lysis buffer and RNA was extracted immediately after sorting.

### RNA isolation and next-generation sequencing

Tissue samples were flash frozen in liquid nitrogen (whole ganglia), or FAC sorted (Fast Blue traced). Then RNA was isolated with the RNeasy Micro Kit (Qiagen) according to the manufacturer’s protocol, including DNase-I digestion. RNA quality was tested via gel electrophoresis (2200 TapeStation, Agilent Technologies) and each sample was assigned an RNA Integrity Number (RIN) based on the RNA quality. Only samples with a RIN > 7 were used for sequencing. RNA-seq was performed on an Illumina HiSeq 2500 at the Duke Center for Genomic and Computational Biology, using single-end sequencing of 50 bp reads. Samples were pooled 4 per lane.

For the whole ganglia LPS sequencing dataset, there were 10 total samples corresponding to 10 female mice (8-10 weeks old); of the 10 mice, 5 were vehicle (healthy control) and 5 were LPS (sick). For the Fast Blue tracing RNA sequencing dataset, n = 2 vehicle, n = 2 LPS. Where Fast Blue positive samples contained only neurons traced from the airways, and the Fast Blue negative samples contain mostly neurons and a fraction of satellite cells. Each sample represents a pool of 5 mice to yield the minimum cell density required for fluorescence-activated cell sorting and high-quality RNA extraction. The sequencing results were aligned to the genome using TopHat (85) and transcripts were mapped to the reference genome, (mm9 which was mapped by the Genome Bioinformatics Group of UC Santa Cruz) using Cufflinks (86). The gene expression values were calculated for each gene in a sample based on the number of Fragments mapped Per Kilobase of transcript per Million reads mapped (FPKM). The number of reads aligned was > 82% for all samples.

Differential expression was determined in two ways, first by using Cuffdiff and a second, manual method, using Matlab with the following criteria: average FPKM > 1, standard error < 33% of the mean, p < 0.05, and at least a 1-fold change up or down. All the resulting genes were further filtered with the following criteria: the higher value (vehicle or LPS) was > 2 FPKM, greater than 2-fold increase for up-regulated genes or greater than 1-fold difference for down-regulated genes. Based on these stringent restrictions only the most strongly regulated genes are presented.

RNA sequencing was loaded into ©Qlucore Omics Explorer (Version 1, Lund Sweden) for Principal Component Analysis (PCA) and heat map generation. PCA was used to visualize the data set in three-dimensional space, after filtering our variables with low overall variance to reduce the impact of noise, and centering and scaling the remaining variables to zero mean and unit variance. The projection score was used to determine the optimal filtering threshold, based on p-value.

### RNA isolation and real time qPCR

Tissue samples (n=4, unless otherwise stated) were surgically removed and snap-frozen by immersion in liquid nitrogen. Total RNA was isolated with the RNeasy Micro Kit (Qiagen, Maryland) according to the manufacturer’s protocol, including DNase-I digestion. RNA quality was tested via gel eletrophoresis (TapeStation, Aligent Technologies) and each sample was assigned an RNA Integrity Number (RIN) according the RNA quality. Only samples with a RIN > 7 were used for qPCR. Concentration was determined using the NanoDrop 1000. For real time qPCR cDNA was made using the High Capacity cDNA Reverse Transcription kit (Applied Biosystems) based on the manufacturer’s protocol. Real time qPCR was performed on a LightCycler 480 (Roche, Indianapolis, IN). The expression of genes of interest was determined using the following TaqMan® Gene Expression Assays from Applied Biosystems (Foster City, CA): Actb (Mm00607939_s1), Lcn2 (Mm01324470_m1), Atf3 (Mm00476032_m1), Sprr1a (Mm01962902_s1), Gpr151 (Mm00808987_s1), Cyp26a1 (Mm00514486_m1), Cckbr (Mm00432329_m1), Npy1r (Mm00650798_g1). The CT value of each well was determined using the LightCycler 480 software and the average of the triplicates was calculated. The relative quantification was calculated as 2-DCt using 18S and Act-b as housekeeping genes (87). All the qPCR data conform to the MIQE guidelines (88).

### Calcium imaging

Calcium imaging was performed 2 hours after ganglia dissociation and culture. Cells were washed once with calcium free PBS then incubated for 45 minutes at 37 °C with 5 µM Fura-2 AM, cell permeant (Life Technologies, Carlsbad, CA) and 0.1% Pluronic F-127 (Life Technologies) in imaging buffer (120 mM NaCl, 3 mM KCl, 2 mM CaCl2, 2 mM MgCl2, 10 mM HEPES, 10 mM glucose). The loading buffer was removed and cells were washed two more times and placed in the dark for 15 minutes to reach room temperature. Radiometric calcium imaging was done using an Olympus IX51 microscope with a Polychrome V monochromator (Till Photonics, Gräfelfing, Germany) light source and a PCO Cooke Sensical QE camera running Imaging Workbench 6 software (INDEC BioSystems, Santa Clara, CA). Fura-2 emission images were obtained with of 50 ms at 340-nm followed by 30 ms at 380-nm excitation wavelengths. Intracellular calcium changes were measured from the F340/F380 ratio. Ratiometric images were generated using ImageJ software (https://imagej.nih.gov/ij/).

### Statistics

Data are given as mean ± SEM. Statistical analyses were carried out using Matlab and Excel. Student’s t-test was used for a single comparison between 2 groups. Results of p-values < 0.05 were considered significant.

## ACKNOWLEDGMENTS

Supported by NIH grant R01HL105635 to SEJ. Flow cytometry was performed at the Duke Human Vaccine Institute Research Flow Cytometry Shared Resource Facility (Durham, NC), which received partial support for construction from the National Institutes of Health, National Institute of Allergy and Infectious Diseases (UC6-AI058607).

## SUPPLEMENTARY MATERIAL

**Figure S1:**
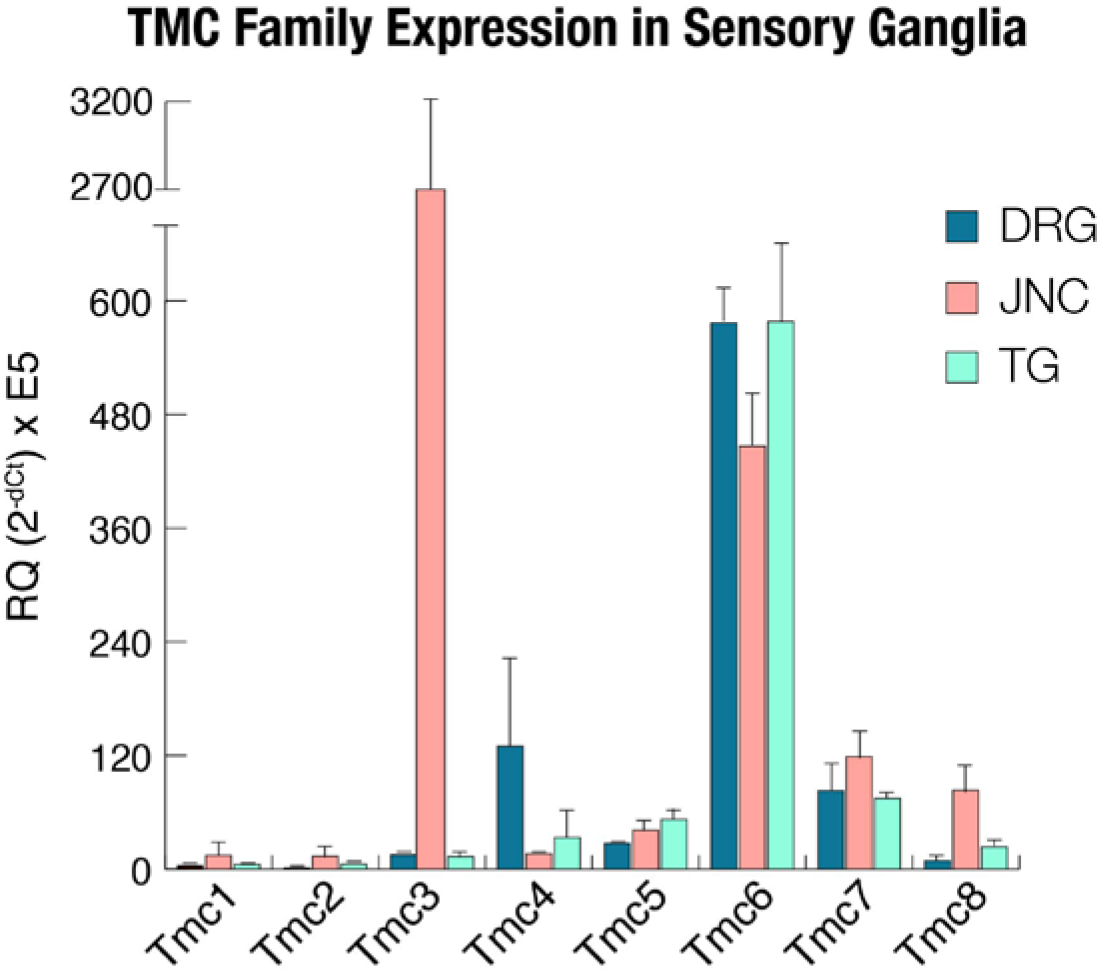
TMC family expression in sensory ganglia. Expression values determined by real time qPCR of Tmc1-8 in DRG, JNC and TG. N=4, values are reported as mean ± SEM.

**Figure S2:**
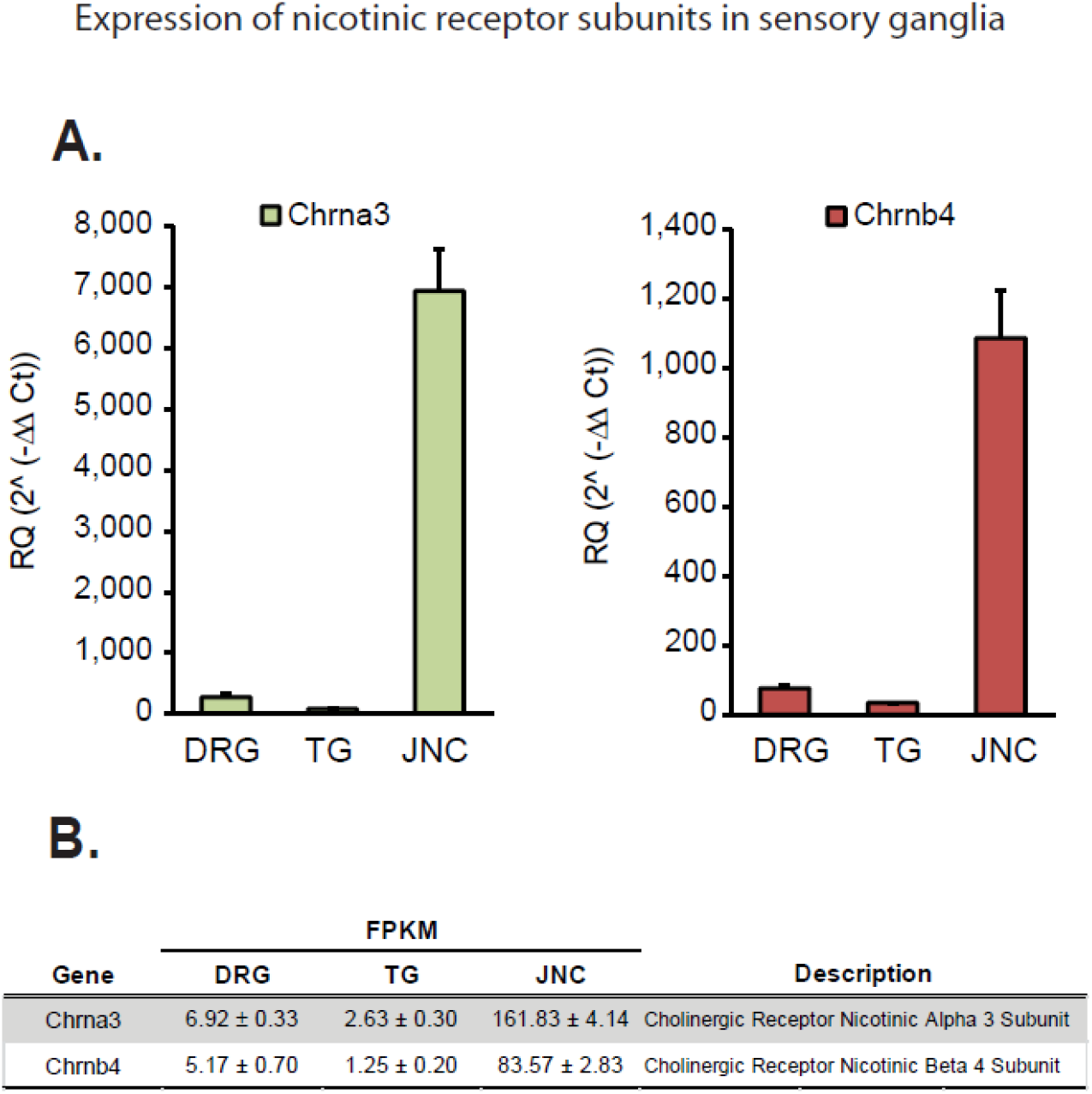
Nicotinic receptor subunits expression in sensory ganglia: **(A)** qPCR expression values of Chrna3 and Chrnb4 subunits in DRG, TG and JNC. N=4, values are reported as mean ± SEM. **(B)** FPKM in DRG, TG and JNC of the same nicotinic receptor subunits.

**Figure S3:**
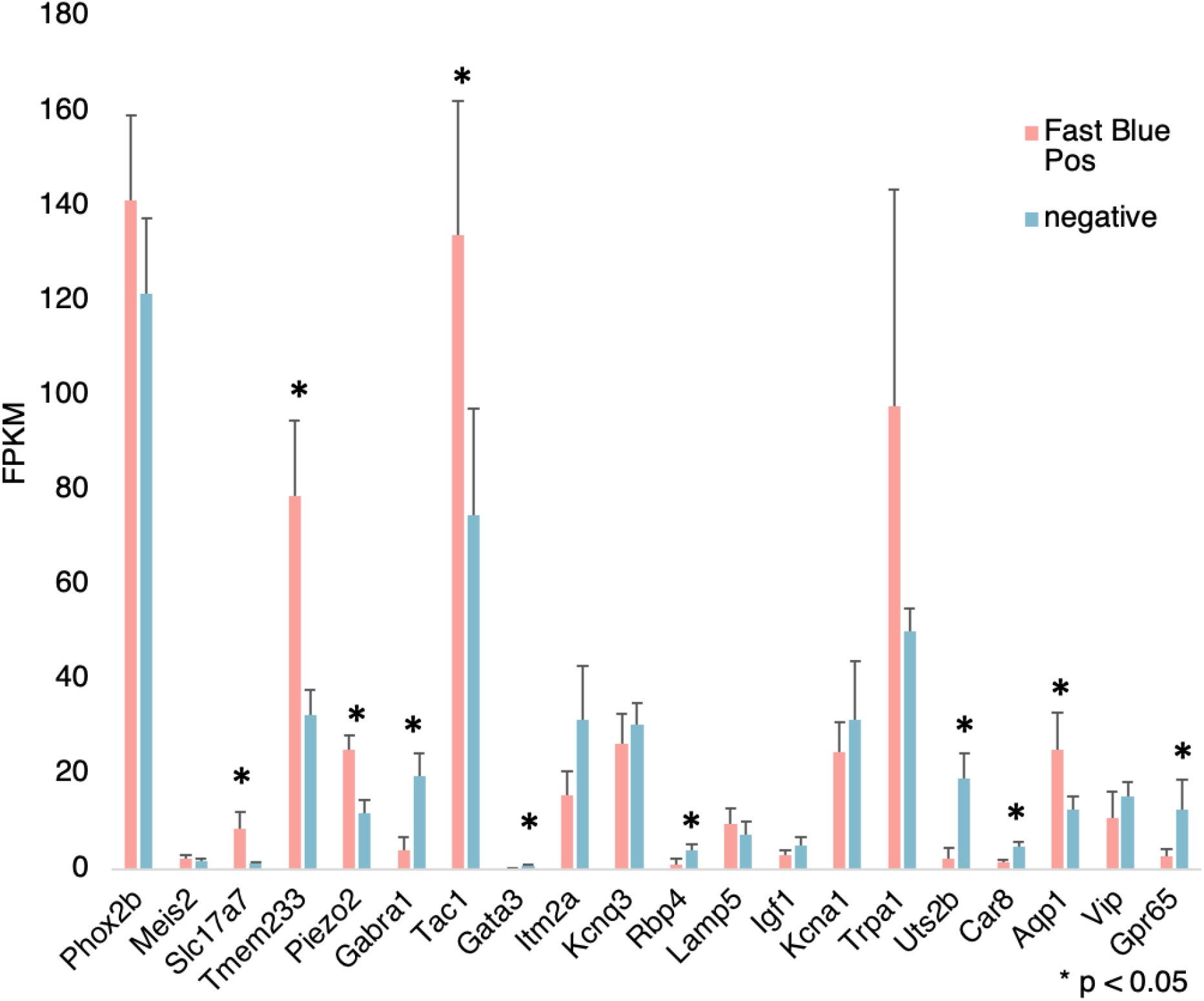
RNAseq FPKM expression of JNC clusters: RNAseq expression values (FPKM) of genes identified to differentially label different nodose ganglia single cell clusters (10) comparing Fast Blue labeled (lung innervating) JNC population to the non-labeled (non lung-innervating) negative population. N=4, values are reported as mean ± SEM; * p < 0.05.

**Table S1:** Alignment Fast Blue positive and negative: RNAseq read alignment and total cell values for the Fast Blue traced populations. The Fast Blue traced neurons represented ∼2% of the total population of neurons in the JNC. Veh = vehicle, LSP = lipopolysaccharide, FB+ = Fast Blue positive JNC neurons, FB-= Fast Blue negative JNC neurons. N=2, each N represents 5 pooled mice.

| Sample | # Cells | Total reads | % reads aligned | % Multiple alignments | # Reads with $\geq 20$ alignments |
| --- | --- | --- | --- | --- | --- |
| Veh1 FB+ | 785 | 29,322,446 | 83.8% | 6.0% | 745 |
| Veh1 FB- | 38,650 | 33,463,139 | 84.3% | 5.3% | 1,566 |
| LPS1 FB+ | 752 | 31,198,754 | 83.3% | 6.9% | 905 |
| LPS1 FB- | 26,266 | 30,231,450 | 85.5% | 5.1% | 1,170 |
| Veh2 FB+ | 1,355 | 30,723,914 | 84.3% | 5.8% | 1,078 |
| Veh2 FB- | 54,963 | 29,977,479 | 85.2% | 3.5% | 2,855 |
| LPS2 FB+ | 1,390 | 32,572,505 | 82.6% | 7.0% | 1,189 |
| LPS2 FB- | 57,212 | 38,033,182 | 87.4% | 4.2% | 2,533 |

**Table S2:** JNC genes Fast Blue positive and negative: RNAseq expression values in FPKM of the 7 highly enriched JNC genes as compared to DRG and TG. WG values represent the whole ganglia expression and is compared to the lung-inntervating neurons (FB+), and the non-lung innervating neurons (FB-). WG = whole ganglia, FB+ = Fast Blue positive JNC neurons, FB-= Fast Blue negative JNC neurons. N=2, each N represents 5 pooled mice, values reported as mean ± SEM.

| Gene | WG | FB+ | FB- |
| --- | --- | --- | --- |
| Phox2b | 90.98 ± 2.29 | 155.83 ± 6.25 | 113.10 ± 2.78 |
| P2rx2 | 90.37 ± 3.04 | 70.91 ± 5.99 | 94.43 ± 26.19 |
| Tmem255b | 28.78 ± 4.00 | 10.07 ± 2.53 | 38.25 ± 11.98 |
| Gpr65 | 26.62 ± 0.42 | 1.91 ± 0.71 | 13.59 ± 5.69 |
| Uts2b | 34.77 ± 5.09 | 1.92 ± 1.73 | 23.24 ± 2.30 |
| Tmc3 | 59.74 ± 7.26 | 36.02 ± 1.49 | 74.60 ± 6.74 |
| Actb | 647.8 ± 31.7 | 679.4 ± 3.41 | 592.52 ± 114.1 |

**Movie S1. Airway-innervating JNC neurons traced with Fast Blue**: Three-dimensional reconstruction of JNC ganglion. Neurons are labeled by Fast Blue that was exposed intranasally one week before collection.

